# Eye movements of younger and older adults decrease during story listening in background noise

**DOI:** 10.1101/2025.04.15.648919

**Authors:** Björn Herrmann, Florian Scharf, Andreas Widmann

## Abstract

Assessments of listening effort are increasingly relevant to understanding the speech-comprehension difficulties experienced by older adults. Pupillometry is the most common tool to assess listening effort but has limitations. Recent research has shown that eye movements decrease when listening is effortful and proposed indicators of eye movements as alternative measures. However, much of the work was conducted in younger adults in trial-based sentence-listening paradigm, during concurrent visual stimulation. The extent to which eye movements index listening effort during continuous speech listening, independently of visual stimuli, and in older adults, is unknown. In the current study, younger and older adults listened to continuous stories with varying degrees of background noise under free and moving-dot viewing conditions. Eye movements decreased (as indexed by fixation duration, gaze dispersion, and saccade rate) with increasing speech masking. The reduction in eye movements did not depend on age group or viewing conditions, indicating that eye movements can be used to assess effects of speech masking in different visual situations and in people of different ages. The pupil area was only sensitive to speech masking early in the experiment. In sum, the current study suggests that eye movements are a potential tool to assess listening effort during continuous speech listening.

## Introduction

Speech comprehension in the presence of background masking sound requires a listener to invest more cognitively, for example, relying more on attention and memory, which makes listening effortful (Eckert et al., 2016; Pichora-Fuller et al., 2016; Peelle, 2018; Herrmann and Johnsrude, 2020a). Assessing listening effort is increasingly relevant in the study of speech-comprehension challenges in older adulthood, because older listeners may experience listening effort in noisy situations long before they are diagnosed with having a hearing loss (Pichora-Fuller and Levitt, 2012; Helfer and Jesse, 2021). Listening effort may thus be a potentially early diagnostic marker of hearing loss.

The most common objective tool to assess listening effort is pupillometry (van der Wel and van Steenbergen, 2018; Winn et al., 2018; Zekveld et al., 2018). The pupil dilates when listening effort increases due to masking of speech (Zekveld et al., 2010; Koelewijn et al., 2018; Zekveld et al., 2019; Cui and Herrmann, 2023; Herrmann and Ryan, 2024), degradation of speech (Zekveld Adriana et al., 2023; Kılıç et al., 2024), or linguistic speech-comprehension challenges (Wendt et al., 2016; Ayasse and Wingfield, 2018; Kadem et al., 2020). However, measuring the pupil also has limitations because its size is sensitive to changes in light (Knapen et al., 2016; Suzuki et al., 2019; Thurman et al., 2021) and the angle of the eye relative to the eye-tracking camera, among other factors (Brisson et al., 2013; Hayes and Petrov, 2016; Fink et al., 2023). To account for the latter, participants are typically instructed to fixate on a point on a computer screen (Ohlenforst et al., 2017; Zekveld et al., 2018; Farahani et al., 2020; Winn and Teece, 2021), but this might reduce external validity because it can reduce memory and mental imagery (Johansson et al., 2012) and reduce behavioral effects of speech comprehension (Cui and Herrmann, 2023). Further, pupillometry is most commonly used in trial-by-trial sentence-listening paradigms (Zekveld et al., 2010; Winn et al., 2015; Wendt et al., 2016; Ayasse and Wingfield, 2018; Borghini and Hazan, 2018; Zekveld et al., 2019; Kadem et al., 2020; Winn and Teece, 2021), whereas an increasing number of works leverage continuous story-listening paradigms to mitigate the less naturalistic nature of short, disconnected sentences (Lalor and Foxe, 2010; Ding and Simon, 2014; Broderick et al., 2018; Brodbeck and Simon, 2020; Panela et al., 2024). The extent to which pupillometry is sensitive to listening challenges under such continuous conditions is a topic of increasing interest (Zhao et al., 2019; Seifi Ala et al., 2020; Fiedler et al., 2021; Cui and Herrmann, 2023; Widmann et al., 2025).

A recent line of studies suggests that eye movements may provide an alternative objective measure to assess listening effort. Eye movements decrease when listening effort increases, for example, due to background noise that masks speech (Contadini-Wright et al., 2023; Cui and Herrmann, 2023; He et al., 2024; Herrmann and Ryan, 2024). Eye movements also decrease during periods of high compared to low memory load (Dalmaso et al., 2017; Kosch et al., 2018; Walter and Bex, 2021) and due to visual-task difficulty (Nakayama et al., 2002), suggesting domain-general reductions in eye movements under high cognitive load. Moreover, eye movements appear to be sensitive to speech masking even during continuous story listening (Cui and Herrmann, 2023). However, in many of the previous works, including the study using continuous story listening (Cui and Herrmann, 2023), some visual stimulus was presented on the computer screen (e.g., fixation point, one moving dot, several moving dots). This visual dot stimulation was used to increase the likelihood of eye movements, allowing for a better investigation of their reduction due to listening effort (Cui and Herrmann, 2023; Herrmann and Ryan, 2024). It is thus unclear whether visual stimulation is advantageous for assessments of listening effort through eye movements or whether free viewing on a blank screen is as effective. Finally, the relationship between eye movements and listening effort has thus far mainly been investigated in younger, normal-hearing adults (Contadini-Wright et al., 2023; Cui and Herrmann, 2023). In a recent sentence-listening study, eye movements were also sensitive to increased speech masking in older adults (Herrmann and Ryan, 2024), but whether this generalizes to continuous story listening is unclear.

In the current study, younger and older adults listened to continuous stories masked by background babble at different signal-to-noise ratios (SNRs) either while a blank screen or a moving-dot display was presented. The aim was to investigate whether pupil area and eye movements are sensitive to speech masking (as a manipulation that induces listening effort) during naturalistic speech listening, and whether the sensitivity to speech masking differs between viewing conditions and/or age groups.

## Methods and Materials

### Participants

Thirty-five younger adults (median age: 23 years, age range: 18–33 years; 16 male, 18 female, 1 non-binary) and thirty-one older adults (median age: 69 years, age range: 55–80 years; 12 male, 19 female) participated in the current study. Participants were native English speakers or learned English before the age of 5 years. They reported having normal hearing and none wore or were prescribed hearing aids. Participants gave written informed consent prior to the experiment and were paid $7.5 CAD per half-hour for their participation. The study was conducted in accordance with the Declaration of Helsinki, the Canadian Tri-Council Policy Statement on Ethical Conduct for Research Involving Humans (TCPS2-2014), and was approved by the Research Ethics Board of the Rotman Research Institute at Baycrest Academy. Data from four additional participants were not analyzed, because recordings in half (N=3) or all (N=1) of the experimental blocks failed due to technical problems.

### Hearing assessment

All experimental procedures were conducted within a sound booth. Pure-tone audiometric thresholds were obtained for 0.25 to 8 kHz using a Maico M28 audiometer. The pure-tone average threshold (PTA) was calculated as the mean threshold across 0.5, 1, 2, and 4 kHz frequencies (Stevens et al., 2013; Humes, 2019). Audiometry was not recorded for two younger and two older participants (administration was simply missed during the recording session). An independent-samples t-test was used to analyze age-group differences in the pure-tone average threshold.

All participants self-reported having normal hearing, but some showed clinically relevant hearing loss (PTA > 20 dB HL; Stevens et al., 2013; Humes, 2019; WHO, 2024; see below). All data were included in the analysis, because the sample is representative of community-dwelling adults (Moore, 2007; Plack, 2014; Presacco et al., 2016; Herrmann et al., 2018, 2023) and the current study aimed to investigate more generally whether eye movements are indicative of listening effort during story listening in younger and older adults. Correlations of eye-movement metrics with PTA assessed the effects of hearing loss as described below.

### Stimulus materials and procedure

Participants listened to four approximately 7.5-min stories from the storytelling podcast The Moth (https://themoth.org/; “Model Magic” by Isobel Connelly [7:30 min]; “Going the Extra Mile” by Luanne Sims [7:36 min]; “The Scare Dee Cats” by Michael Donovan [7:18 min]; “The Long Way Home” by Tod Kelly [7:45 min]). The Moth is a podcast where people tell stories about interesting life events. Stories are highly enjoyable and absorbing (Herrmann and Johnsrude, 2020b; Irsik et al., 2022b; Cui and Herrmann, 2023). Each participant listened to two stories while a blank gray screen was presented, whereas they listened to the other two stories while 16 white dots moved randomly on the screen, as described in detail previously (Cui and Herrmann, 2023; Herrmann and Ryan, 2024), mirroring the stimulation of multiple object tracking paradigms. (Cavanagh and Alvarez, 2005; Alvarez and Franconeri, 2007; Scholl, 2009; Herrmann and Johnsrude, 2018b, a). The moving-dots display aimed to facilitate eye movements (see Cui and Herrmann, 2023), but no task was required to avoid a dual-task procedure. Participants were instructed to look at the screen in whatever way they wanted (Johansson et al., 2006; Johansson et al., 2011; Johansson et al., 2012; Herrmann and Ryan, 2024). The viewing conditions were blocked such that both stories of the same viewing condition were presented in direct succession (starting condition was counterbalanced across participants). The assignment of stories to the viewing conditions and the order in which stories were presented was counterbalanced across participants.

Each story was masked by 12-talker background babble (Bilger, 1984; Wilson et al., 2012). The SNR between the speech signal and the 12-talker babble changed every 27 seconds to one of five SNR levels (+17, +12, +7, +2, -3 dB SNR; for a similar approach see Irsik et al., 2022b, a; Cui and Herrmann, 2023), corresponding to about a range of 95% to 55% of intelligible words (Irsik et al., 2022b). Each SNR was manipulated by adjusting the dB level of both the story and the masker. This ensured that the overall sound level remained constant across SNRs and throughout a story, and that the overall level was similar for both stories. Each story started and ended with the +17-dB SNR level to enable participants to clearly hear the beginning and end of the story (these were excluded from the analysis, see below). Each SNR level was presented 3 times in a story, with the exception of the +17 dB SNR level, which was presented 5 times (3 times + beginning and end). The SNR transitioned smoothly from one level to the other over a duration of 1 second. The order of SNR levels was randomized such that a particular SNR could not be heard twice in succession, and that SNR would maximally change by two levels. For each participant, SNR levels were randomized uniquely.

The four stories were presented in four separate blocks, and participants took a break between blocks (median: 5 min; mean: 6 min; range: 2–25 min). Each story was presented continuously for the ∼7.5 min duration without any silence periods or interruptions (SNR levels seamlessly transitioned; Irsik et al., 2022b, a; Cui and Herrmann, 2023). The naturalistic story-listening paradigm thus did not contain any silent or non-speech baseline periods nor unique disconnected trials. After a story ended, participants answered ten comprehension questions about the story. For each comprehension question, participants were asked to select the correct answer out of four multiple choice options.

Prior to the main experiment procedures, participants underwent a familiarization block in which they listened to a short 1:12 min story.

### Experimental setup

Sounds were presented via Sony Dynamic Stereo MDR-7506 headphones and a Steinberg UR22 mkII (Steinberg Media Technologies) external sound card. Experimental procedures were run using Psychtoolbox (v3.0.14) in MATLAB (MathWorks Inc.) on a Lenovo T450s laptop with Microsoft Windows 7. The laptop screen was mirrored to a ViewSonic monitor with a refresh rate of 60 Hz. All sounds were presented at a comfortable listening level that was fixed across participants (∼70-75 dB SPL).

### Behavioral data analysis

The proportion of correct responses was calculated for the story comprehension task, separately for each story. The proportion of correct responses were averaged across the two stories for each viewing condition (blank screen, moving-dot display). A repeated measures analysis of variance (rmANOVA) was calculated with the within-participants factor Viewing Condition (blank, dots) and the between-participants factor Age Group (younger, older).

### Pupillometry and eye-movement recordings

During the experiments, participants placed their head on a chin and forehead rest facing the computer monitor at a distance of about 70 cm. Pupil area and eye movements were recorded continuously from the right eye (or the left eye if the right eye could not be tracked accurately) using an integrated infrared camera (EyeLink 1000 eye tracker; SR Research Ltd.) at a sampling rate of 500 Hz (centroid tracking mode). Nine-point fixation was used for eye-tracker calibration prior to each block (McIntire et al., 2014).

### Processing of pupil-area and eye-movement data

The main metrics of the current study were pupil size, fixation duration, gaze dispersion, and (micro-) saccade rate (Cui and Herrmann, 2023; Herrmann and Ryan, 2024). Fixation duration and gaze dispersion are broad measures of eye movements that we recently used to non-specifically capture any eye movement changes associated with listening effort (Cui and Herrmann, 2023; Herrmann and Ryan, 2024). A broad measure of eye movements may be particularly advantageous if an index of listening effort is the focus rather than understanding underlying mechanisms. Nevertheless, we also analyzed (micro-) saccade rate (Engbert and Kliegl, 2003; Engbert, 2006; Widmann et al., 2014) since it can be sensitive to memory load and listening effort in trial-based tasks (Dalmaso et al., 2017; Contadini-Wright et al., 2023; Kadosh et al., 2024), although this is not always found (Kadem et al., 2020; Cui and Herrmann, 2023). Henceforth, we refer to this measure simply as saccade rate, acknowledging the fact that micro-saccades and saccades likely share functional characteristics (Otero-Millan et al., 2008; Martinez-Conde et al., 2009; Martinez-Conde et al., 2013).

Preprocessing was calculated for each experimental block separately using MATLAB. For each eye blink indicated by the eye tracker, all data points of the pupil-area time course and the x and y eye-coordinate time courses between 100 ms before and 200 ms after a blink were set to NaN (‘not a number’ in MATLAB). Moreover, pupil-area values that differed from the mean pupil area (across a block) by more than 3 times the standard deviation were classified as outliers and the corresponding data points of the pupil-area time course and the x and y eye-coordinate time courses were set to NaN. MATLAB’s ‘pchip’ method was used to interpolate NaN-coded samples in the pupil data. Pupil-size time courses were filtered with a 5-Hz low-pass filter (FIR, 51 points, Kaiser window, β = 4). X and y eye-coordinate time courses were not interpolated. That is, missing data points (NaNs) were ignored in the calculation of the eye-movement metrics.

Broad changes in eye movements were investigated using fixation duration and gaze dispersion, similar to our previous work (Cui and Herrmann, 2023; Herrmann and Ryan, 2024). Both measure the general tendency for the eyes to move around. Fixation duration was calculated as the time a person’s x-y eye coordinates remained in a given location (within 0.5° visual angle; radius of 10 px). For each time point, the corresponding x-y coordinate defined the center of the critical 0.5° visual-angle location. The number of continuous pre- and post-samples was calculated for which the x-y-eye coordinates remained in the defined location. The sample number was divided by the sampling frequency to obtain the fixation duration for the specific time point. If a data value of any pre- or post-sample within the 0.5° visual angle location had been coded NaN (i.e., was missing), the fixation duration of the corresponding time point was set to NaN and ignored during averaging.

Gaze dispersion was calculated as the standard deviation in gaze across time points, averaged across x- and y-coordinates, and transformed to logarithmic values to make the metric’s distributional properties more Gaussian. Smaller values indicate less gaze dispersion. To obtain time courses for gaze dispersion, it was calculated for 1-s sliding time windows centered sequentially on each time point. If more than 90% of data were unavailable within a 1-s time window (that is, 450 or more samples were NaN-coded), gaze dispersion for the corresponding time point was set to NaN and ignored during averaging (Cui and Herrmann, 2023; Herrmann and Ryan, 2024).

Saccade rate was calculated as a more specific measure of eye movements. Saccades/ microsaccades were identified using a method that computes thresholds based on velocity statistics from x- and y-coordinate trial time courses and then identifies saccades/microsaccades as events passing that threshold (Engbert and Kliegl, 2003; Engbert, 2006). Similar to gaze dispersion, a 1-s sliding window was sequentially moved across the data of an experimental block. For each 1-s time window, the vertical and horizontal eye movement time series were transformed into velocities, and saccades/mircosaccades were classified as outliers if they exceeded a relative velocity threshold of 15 the standard deviation of the eye-movement velocity and persisted for 6 ms or longer. Previous work differed in the specific velocity threshold that was used, with some work using a velocity threshold of 5 (Engbert and Kliegl, 2003; Widmann et al., 2014), others used a threshold of 15 (Kadem et al., 2020), whereas yet other tested several (Cui and Herrmann, 2023). A threshold of 15 was chosen here, because a threshold of 5 lead to an unrealistically high saccade rate (∼10 Hz). If any of the x- or y-values in the eye-tracking time courses comprised an NaN, the saccade rate for this specific sliding window was set to NaN (note that the saccade rate calculation requires continuous data points within the analysis window, prohibiting the presence of NaNs).

### Statistical analysis of pupil area, fixation duration, and gaze dispersion

Pupil area, fixation duration, gaze dispersion, and saccade rate were separately averaged within each 27-s SNR segment of a story. Data from the +17 dB SNR segments at the beginning and end were not used for analysis, because they might otherwise give more weight to this condition in the analysis. For each participant and eye-tracking metric (pupil area, fixation duration, gaze dispersion, saccade rate), this resulted in 15 data points per block, consisting of three times the 5 SNR conditions (+17, +12, +7, +2, -3 dB).

A multi-level regression analysis was performed in MATLAB (Holmes and Friston, 1998; Worsley et al., 2002; Penny and Holmes, 2007; Lindquist, 2008). On the first level, a linear model per eye-tracking metric (i.e., pupil area, fixation duration, gaze dispersion, saccade rate) was fitted separately for each participant. The predictors of interest in the model were SNR (coded: -3, +2, +7, +12, +17 dB, and then zero-centered), Viewing Condition (coded: -0.5, 0.5 for blank and moving dots, respectively), the SNR × Viewing Condition interaction, and an intercept (capturing the overall magnitude). Previous work suggests that modeling time-on-task is important to account for longer term trends in the data (Unsworth and Robison, 2016; Benwell et al., 2018; Fink et al., 2023; McLaughlin et al., 2023; Widmann et al., 2025). Hence, segment number was also included as a linear trend (coded -7 to 7 in increments of 1) and a quadratic trend (squared linear predictor) to account for changes within a block of story listening (i.e., time-on-task). For each participant and eye-tracking metric, the result of the first-level analysis was a coefficient for SNR, Viewing Condition, SNR × Viewing Condition, and the intercept as well as for the linear and quadratic time-on-task trends, describing the relationship with the eye-tracking metric.

On the second level, coefficients were tested against zero using a one-sample t-test (except for the intercept), essentially testing the main effects of SNR and Viewing Condition, and the SNR × Viewing Condition interaction. One-sample t-tests were also conducted for linear and quadratic time-on-task trends, showing significant effects for all four eye-tracking metrics (for all t > 2, p < 0.05; except for the quadratic trend for saccade rate p > 0.1). Linear and quadratic trends were removed for plotting the data and not further considered (e.g., all predicted values depicted in the figures were computed accounting for linear and quadratic trends by fixing the segment at zero). An independent samples t-test on the intercept was calculated to assess the main effect of Age Group. Independent samples t-tests on the coefficients for SNR, Viewing Condition, and SNR × Viewing Condition were also calculated to assess interactions with Age Group. Effect sizes are provided as Cohen’s d (Cohen, 1988).

In an explorative multi-level regression analysis, we also examined whether an SNR effect on eye metrics may be greater in the beginning of the experimental session. To this end, the regressor for Viewing Condition was replaced by a regressor coding for the block number. The experimental design and available data did not allow running a larger regression model including Viewing Condition and all interactions with Block. Analysis of block-wise effects were calculated post hoc and should be considered exploratory.

Additional analyses were calculated to investigate the relationship between changes in the pupil area and changes in the eye-movement metrics (gaze dispersion, fixation duration, saccade rate). To this end, between-participant partial correlations were calculated separately between the pupil area and the three eye-movement metrics using the linear coefficient for the SNR effect (partialling out age group).

We also investigated the between-participant relationship between pure-tone average threshold (PTA) and eye-tracking metrics (i.e., pupil area, fixation duration, gaze dispersion, saccade rate). To this end, separately for each eye-tracking metric, a regression model was calculated using the estimated linear coefficient for SNR from each participant’s linear model as the dependent measure (i.e., the SNR effect) and pure-tone average threshold (PTA) as the predictor. Age group was included as an additional regressor to account for effects of age independent of hearing loss. Additional regressions were calculated to examine whether PTA predicts the overall magnitude of the eye-tracking metrics (using the estimated intercept from each participant’s linear model), again including age group as a regressor.

## Results

Pure-tone average thresholds (PTAs) were greater for older compared to younger adults (t_60_ = 7.163, p = 1.3 × 10^−9^, d = 1.820; Figures 1A and B), as expected based on the known elevated thresholds in community-dwelling older adults (Cruickshanks et al., 1998; Moore, 2007; Plack, 2014; Goman and Lin, 2016; Presacco et al., 2016; Herrmann et al., 2018, 2023). Some older adults had PTAs greater than 20 dB HL, which would be considered clinical hearing loss (Stevens et al., 2013; Humes, 2019; WHO, 2024), but the distribution of PTAs is continuous, suggesting a regression model is appropriate to investigate the impact of hearing function on eye-tracking metrics (see below).

**Figure 1:**
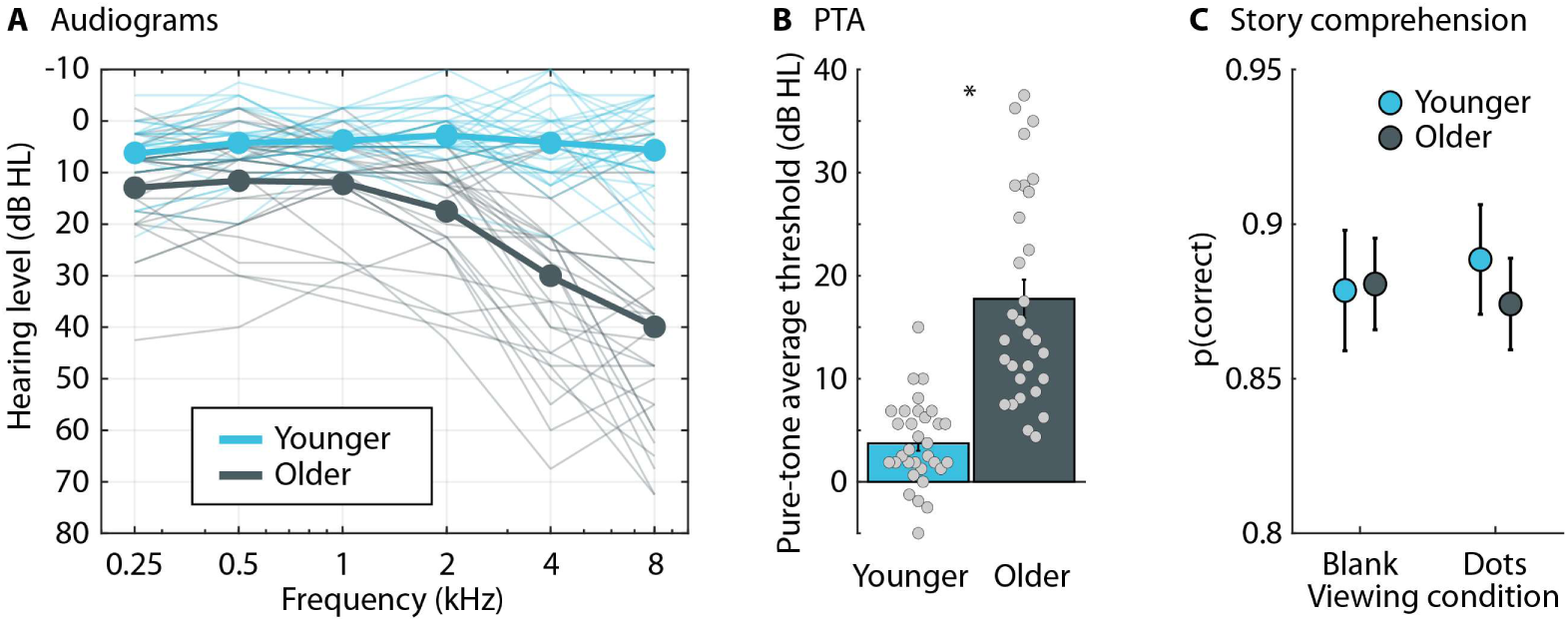
Audiometric and behavioral measures. A: Audiograms. The thin lines show data of individual participants, whereas the thick lines show the mean across participants. B: Pure-tone average thresholds (average across audiometric thresholds at 0.5, 1, 2, 4 kHz). The bar graphs show the mean across participants and individual dots reflect thresholds from individual participants. Error bars reflect the standard error of the mean (SEM). C: Mean and SEM for the story-comprehension assessment. *p < 0.05

**Figure 2:**
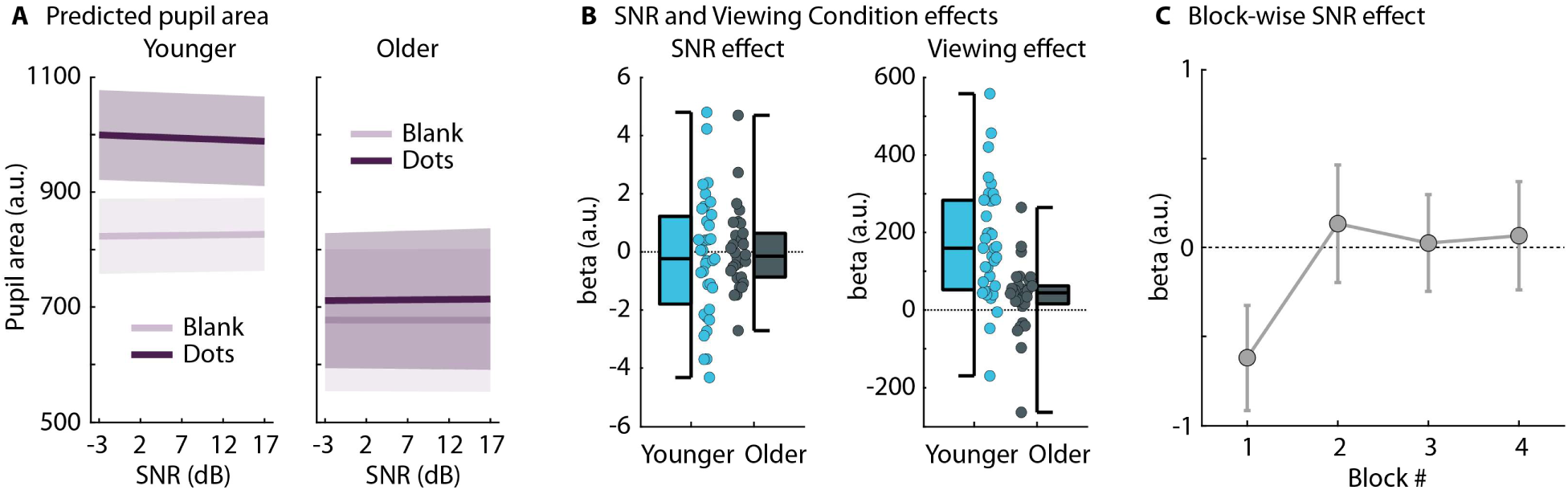
Results for the pupil area. A: Predicted pupil area from fitting a linear model for each participant. The data shown reflect the residuals after removing time-on-task linear and quadratic trends. B: The effects of SNR and viewing condition (blank screen vs moving-dots display). The data reflect the slopes (beta values) from the linear model fits (dots are the slopes from different participants). For the SNR effect, a beta value greater than 0 indicates that the pupil area increased with increasing SNR. For the viewing-condition effect, a beta value greater than 0 indicates that the pupil area was greater for the moving-dots display than the blank screen. C: Mean beta values (slopes) for the SNR effect from an exploratory analysis, separately for each block of the experiment. The SNR effect was significant only for the first block.

Despite elevated thresholds, story comprehension did not differ between age groups (F_1,64_ = 0.087, p = 0.769, ω^2^ < 0.001) nor between the two viewing conditions (F_1,64_ = 0.021, p = 0.886, ω^2^ < 0.001). There was also no interaction (F_1,64_ = 0.447, p = 0.506, ω^2^ < 0.001).

For the analysis of the pupil area, one person’s dataset was removed, because their data point for the Viewing Condition effect (estimated coefficient) was over 7 times the standard deviation below the sample mean. Inclusion vs exclusion of this person’s data had no qualitative impact – in terms of significance – on any of the other effects; none of the conclusions are affected because of the data removal. Multi-level regression modeling to predict the pupil area revealed no effect of SNR (t_64_ = – 0.322, p = 0.749, d = 0.040), and there was no interaction involving SNR (for all t < 1.9, p > 0.05). The pupil area was greater for the moving-dot display than the blank screen (effect of Viewing Condition: t_64_ = 6.130, p = 6.1 × 10^−8^, d = 0.760), and this interacted with Age Group (t_63_ = 4.297, p = 6.1 × 10^−5^, d = 1.069). The larger pupil area for the moving-dot display than the blank screen was significant for both age groups (for both t > 2.1, p < 0.04), but the difference was greater in younger adults. The exploratory analysis to examine differences in the SNR effect across blocks showed a significant SNR × Block interaction (t_64_ = 2.089, p = 0.041, d = 0.259), resulting from an increase in the pupil area as SNR decreased but only in the first block (t_64_ = –2.105, p = 0.039, d = 0.261; no Age Group difference: p > 0.1) and not in any of the other three blocks (for all > 0.6).

The multi-level regression model shows that fixation duration increased as the SNR decreased (more speech masking; effect of SNR: t_65_ = –3.194, p = 0.002, d = 0.393) and was greater for the blank screen than the moving-dots display (effect of Viewing Condition: t_65_ = –2.714, p = 0.009, d = 0.334; Figure 3). Older adults moved their eyes overall more than younger adults (effect of Age Group: t_64_ = 3.576, p = 6.7 × 10^−4^, d = 0.882). There were no interactions (for all t < 1.3, p > 0.2). The exploratory analysis did not reveal a difference in the SNR effect between blocks (SNR × Block interaction: t_65_ = 0.643, p = 0.523, d = 0.079; SNR effects for blocks 1-4: p = 0.004, p = 0.036; p = 0.141, p = 0.148, respectively).

**Figure 3:**
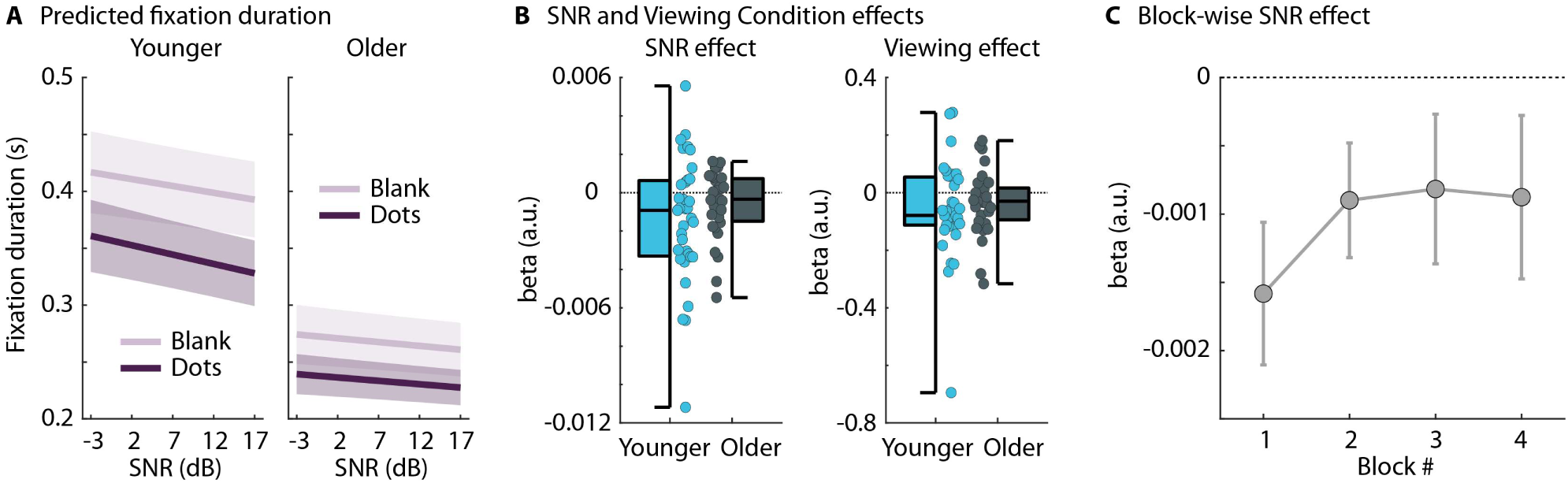
Results for the fixation duration. A: Predicted fixation durations from fitting a linear model for each participant. The data shown reflect the residuals after removing time-on-task linear and quadratic trends. B: The effects of SNR and viewing condition (blank screen vs moving-dots display). The data reflect the slopes (beta values) from the linear model fits (dots are the slopes from different participants). For the SNR effect, a beta value smaller than 0 indicates that fixation durations decreased (i.e., more eye movements) with increasing SNR. For the viewing-condition effect, a beta value smaller than 0 indicates that fixations durations were shorter for the moving-dots display than the blank screen. C: Mean beta values (slopes) for the SNR effect from an exploratory analysis, separately for each block of the experiment. The SNR effect was significant for the first two blocks.

Gaze dispersion mirrored the results from the fixation duration analysis. Multi-level regression modeling revealed a decrease in gaze dispersion as the SNR decreased (more speech masking; effect of SNR: t_65_ = 4.821, p = 9 × 10^−6^, d = 0.593) and gaze dispersion was greater for the moving-dots display than the blank screen (effect of Viewing Condition: t_65_ = 3.509, p = 8.2 × 10^−4^, d = 0.432; Figure 4). Numerically, older adults moved their eyes more than younger adults, but this was not significant for gaze dispersion (effect of Age Group: t_64_ = 1.602, p = 0.114, d = 0.395). There were no interactions (for all t < 0.8, p > 0.4). The exploratory analysis did not reveal any difference in the SNR effect across blocks (SNR × Block interaction: t_65_ = 1.061, p = 0.293, d = 0.131; SNR effects for blocks 1-4: p < 0.001, p = 0.005; p < 0.001, p = 0.005, respectively).

**Figure 4:**
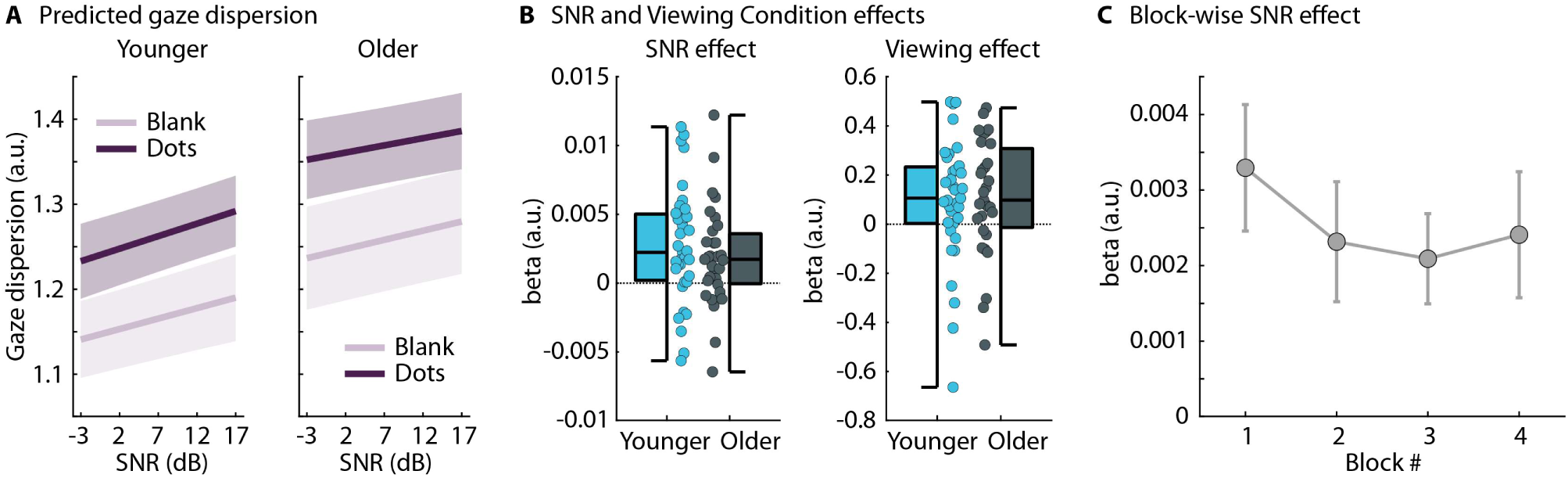
Results for gaze dispersion. A: Predicted gaze dispersion from fitting a linear model for each participant. The data shown reflect the residuals after removing time-on-task linear and quadratic trends. B: The effects of SNR and viewing condition (blank screen vs moving-dots display). The data reflect the slopes (beta values) from the linear model fits (dots are the slopes from different participants). For the SNR effect, a beta value greater than 0 indicates that gaze dispersion increased (i.e., more eye movements) with increasing SNR. For the viewing-condition effect, a beta value greater than 0 indicates that gaze dispersion was greater for the moving-dots display than the blank screen. C: Mean beta values (slopes) for the SNR effect from an exploratory analysis, separately for each block of the experiment. The SNR effect was significant for all blocks.

The analysis of saccade rate also showed comparable results. Participants made fewer saccades/microsaccades as the SNR decreased (more speech masking; effect of SNR: t_65_ = 3.825, p = 3 × 10^−4^, d = 0.471) and for the blank screen than the moving-dots display (effect of Viewing Condition: t_65_ = 5.236, p = 1.9 × 10^−6^, d = 0.645; Figure 5). The saccade rate was overall greater for older than younger adults (effect of Age Group: t_64_ = 3.485, p = 8.9 × 10^−4^, d = 0.86). There were no interactions (for all t < 1.1, p > 0.25). The exploratory analysis did not reveal a difference in the SNR effect between blocks (SNR × Block interaction: t_65_ = 1.119, p = 0.267, d = 0.138; SNR effects for blocks 1-4: p = 0.001, p = 0.115; p = 0.56, p = 0.585, respectively). Note, however, that the estimation of SNR effects for separate blocks was a bit more poorly (Figure 5C), because data points for saccade calculations were sparser.

**Figure 5:**
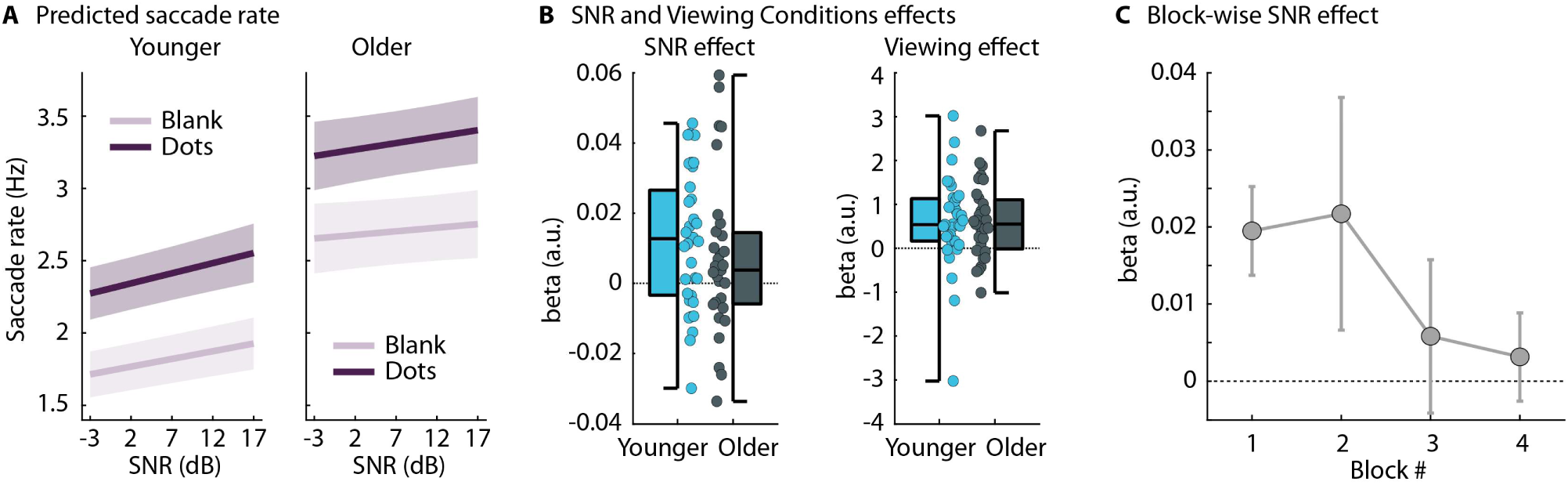
Results for saccade rate. A: Predicted saccade rate from fitting a linear model for each participant. The data shown reflect the residuals after removing time-on-task linear and quadratic trends. B: The effects of SNR and viewing condition (blank screen vs moving-dots display). The data reflect the slopes (beta values) from the linear model fits (dots are the slopes from different participants). For the SNR effect, a beta value greater than 0 indicates that saccade rate increased (i.e., more eye movements) with increasing SNR. For the viewing-condition effect, a beta value greater than 0 indicates that saccade rate was greater for the moving-dots display than the blank screen. C: Mean beta values (slopes) for the SNR effect from an exploratory analysis, separately for each block of the experiment. The SNR effect was significant only for the first block; note though that block-wise estimation of the SNR effect for saccade rate was more poorly (data points are sparser for saccades).

Between-participant correlations between the pupil area and the eye-movement metrics (gaze dispersion, fixation duration, saccade rate) for the linear SNR coefficient revealed no significant relationships (fixation duration: r = 0.215, p = 0.089; gaze dispersion: r = –0.049, p = 0.700; saccade rate: r = –0.117, p = 0.358; partialling out age group). Limiting the between-participant correlations to the SNR effect in the first block of the experiment revealed a significant negative correlation between pupil area and saccade rate (r = –0.253, p = 0.043), whereas the correlation between pupil area and the other two eye-movement measures were not significant (fixation duration: r = 0.111, p = 0.383; gaze dispersion: r = –0.170, p = 0.179). Please note that these correlations are exploratory and should be interpreted with caution.

The SNR effect correlated between the three eye-movement metrics (fixation duration with gaze dispersion: r = –0.638, p = 1.4 × 10^−8^; gaze dispersion with saccade rate: r = 0.765, p = 1.8 × 10^−13^; fixation duration with saccade rate: r = –0.502, p = 2.4 × 10^−5^; partialling out age group).

Finally, regression models were calculated to examine whether the pure-tone average threshold (PTA) predicts SNR-related changes or overall responses of eye-tracking metrics (pupil area, fixation duration, gaze dispersion, saccade rate). For none of the regression did PTA predict SNR-related changes or overall responses of the eye-tracking metrics (for all p > 0.05; also when limited to the 1^st^ block). Older adults moved their eyes more than younger adults as indicated by the significant age-group intercept effects for fixation duration, gaze dispersion, and saccade rate (fixation duration: t_61_ = -2.310, p = 0.024; gaze dispersion: t_61_ = 2.009, p = 0.049; saccade rate: t_61_ = 2.509, p = 0.015; pupil area showed a marginally significant reduction in older adults: t_60_ = -1.971, p = 0.053), which is consistent with the analyses reported above. There were no age group effects for the SNR-related changes in eye-tracking metrics (for all p > 0.7), again consistent with the analyses reported above.

There was also no effect of gender on the overall amplitude nor an interaction with the SNR effect for any of the eye-tracking metrics (for all p > 0.35).

## Discussion

The current study investigated the extent to which eye movements can be used to measure listening effort during story listening in younger and older adults. Participants listened to stories with varying degrees of background masking babble under different viewing conditions (blank screen, moving-dots display). The data show that eye movements decrease when speech masking and, in turn, listening effort increase. The pupil area seemed not very sensitive to speech masking under continuous story listening. The reduction in eye movements under speech masking did not significantly differ between viewing conditions nor age groups, suggesting that eye movements could be used as an effective way to assess listening challenges.

### Effects of listening effort on pupil area decline fast during story listening

Pupil size has repeatedly been shown to increase when listening effort increases in trial-by-trial, sentence-listening paradigms (Zekveld et al., 2010; Kuchinsky et al., 2013; Zekveld et al., 2014; Zekveld and Kramer, 2014; Winn et al., 2015; Wendt et al., 2016; Winn, 2016; Wendt et al., 2017; Zekveld et al., 2019; Kadem et al., 2020; Winn and Teece, 2021; Zhang et al., 2022; Neagu et al., 2023). Other work using ∼30-s speech snippets under different listening challenges found that pupil size can be sensitive to listening effort for longer speech segments, although these works treated pupil data similar to trial-by-trial paradigms (Seifi Ala et al., 2020; Fiedler et al., 2021; for non-speech stimuli see Zhao et al., 2019). Very few studies have used pupillometry to assess listening effort for continuous speech. In the current study, the pupil area was not sensitive to speech masking – a manipulation to induce listening effort – during story listening when the whole dataset was used in the analysis. An explorative block-wise analysis showed that pupil area increased with increasing speech masking, but only in the first experimental block. This is consistent with previous work using a story-listening paradigm that was comparable to the current study, observing that the pupil area is sensitive to speech masking when time-on-task trends are accounted for (Widmann et al., 2025). The speech masking effect in this previous study was also driven mainly by earlier parts of the story, although the effect also came out in the overall analysis (Widmann et al., 2025), whereas this was not the case here.

Potential differences between studies are that an older eye tracker was used for the current than the previous story-listening work (Cui and Herrmann, 2023; Widmann et al., 2025), potentially reducing sensitivity. Perhaps more critical, the overall pupil area appeared smaller in the current study (pupil area of 700-900; Figure 2) than in the previous work (pupil area of 1250-1750; Cui and Herrmann, 2023; Widmann et al., 2025). Although there were no obvious differences in the ambient light conditions of the environments in which the data for these studies were recorded, the smaller pupil area might indicate that more light was reaching participants’ eyes in the current than the previous work. Recommendations for recording pupillometry for assessments of listening effort favor brighter over darker ambient light conditions (Winn et al., 2018), and a relatively small overall pupil area in the current study would suggest that there is the capacity for the pupil area to increase under listening challenges. However, effects of arousal on the pupil size have been reported to depend on light level and may even be abolished at high retinal illuminance (i.e., small pupil size; Pan et al., 2024). It may thus be that retinal illuminance impacted the effect of listening effort on pupil size. A few works have suggested that individual adjustments of the ambient light level may be beneficial for assessing listening effort through pupillometry (Zekveld et al., 2010; Koelewijn et al., 2014; Ohlenforst et al., 2017; Zekveld et al., 2019). Nevertheless, another previous study from our lab, using exactly the same setup and environmental conditions as the current study, observed pupil increases during sentence listening (Herrmann and Ryan, 2024), thus indicating that sensitivity to speech masking per se is not reduced in the current setup.

Regardless of the reasons of why the pupil area may not have been as sensitive to SNR in the current than previous work, the collective data may suggest that using pupil size to assess listening effort during story listening may depend on parameters of the eye tracker, recording environment, and time-on-task, and may not be as easily observed as during trial-by-trial, sentence-listening paradigms. Optimizing the setup for pupil recordings in future work and accounting for time-on-task (Widmann et al., 2025) may thus be critical for capturing effects of SNR on the pupil area during story listening.

### Eye movements provide a window onto listening effort

The current study shows that the number and distance of eye movements decrease (indexed by gaze dispersion, fixation duration, and saccade rate) when the level of background noise relative to speech increases (Figures 3-5), consistent with a few other recent works (Contadini-Wright et al., 2023; Cui and Herrmann, 2023; He et al., 2024; Herrmann and Ryan, 2024). Eye movements may thus indicate the degree of listening effort.

Critically, the current study shows that eye movements decrease with increasing speech masking during story listening for both the free-viewing (blank screen) and moving-dots conditions. Previous studies presented a blank screen, a single dot, or a fixation cross during sentence listening or several dots during sentence or story listening (Contadini-Wright et al., 2023; Cui and Herrmann, 2023; Herrmann and Ryan, 2024). These studies jointly with the current data indicate that the reduction in eye movements during cognitively challenging listening generalizes across visual-stimulation contexts. Eye movements may thus be an effective tool, especially in situations where visual information or light conditions cannot be controlled well and where, in turn, pupillometry may be less feasible.

Older adults moved their eyes more overall (Figures 3, 5), which is a well-known phenomenon (Ryan et al., 2007; Liu et al., 2018; Mazloum-Farzaghi et al., 2022), although the reasons for this are less clear. Critically, age group did not modulate the SNR effect, indicating that eye movement reductions due to listening challenges were mostly similar in younger and older adults. Moreover, the pure-tone average threshold, indexing the degree of a person’s peripheral hearing loss (at least outer hair cell loss; Oxenham and Bacon, 2003; Moore, 2007), also did not modulate the effect of SNR on eye movements. The similar modulation of eye movements by SNR across age groups and PTAs may be somewhat surprising, given that older adults consistently show lower speech intelligibility in sentence-listening paradigms for given SNRs (Helfer and Freyman, 2008; Ferguson et al., 2010; Presacco et al., 2019; Sobon et al., 2019; Pandey and Herrmann, in press). However, younger and older people are similarly absorbed by naturalistic stories (Mathiesen et al., 2024) and story absorption appears to be little impacted by moderate background noise (Herrmann and Johnsrude, 2020b). Older adults have also been shown to benefit well from speech context (Payne and Silcox, 2019) and tend to report as many details and comparable gist comprehension in more naturalistic, continuous listening paradigms (Gordon et al., 2009). The latter is consistent with the current behavioral results showing that both younger and older adults have similarly high story-comprehension accuracy (∼0.87). The current data thus provide little evidence that older adults experienced more listening effort than younger adults, but that eye movements are sensitive within participants to the listening challenges associated with speech masked by background babble.

### Potential mechanisms of reduced eye movements

In the current study, we employed different analyses to capture changes in eye movements. Fixation duration and gaze dispersion have recently been used as broad measures of eye movements – that can include saccades and smooth pursuit among others – to obtain an all-encompassing measure of listening effort (Cui and Herrmann, 2023; Herrmann and Ryan, 2024). Here we also used a saccade/microsaccade rate as a more specific measure (Engbert and Kliegl, 2003; Engbert, 2006; Widmann et al., 2014), which has shown sensitivity to cognitive demands in previous studies (Dalmaso et al., 2017; Contadini-Wright et al., 2023; Kadosh et al., 2024), although not all (Kadem et al., 2020; see also Cui and Herrmann, 2023). The pattern of results of the three eye-movement metrics were very similar and correlated, suggesting that any of the metrics could be used. Nevertheless, SNR effects of gaze dispersion were significant for all experimental blocks, potentially pointing to better sensitivity (although rather numerically in the current study), especially when indexing listening effort is the main purpose of a study.

The effects of speech masking differed somewhat between the pupil area and eye-movement metrics, although it is unclear whether this reflects differences in the measures’ sensitivity or different underlying mechanisms. Cortical and subcortical brain structures – such as the visual cortex, prefrontal cortex, posterior parietal cortex, frontal and supplementary eye fields, anterior cingulate cortex, cerebellum, thalamus, basal ganglia, and superior colliculus – underlie the generation and regulation of eye movements (Sparks, 2002; Pierrot-Deseilligny et al., 2004; Pierce et al., 2019). Changes in the pupil size are driven by some of the same regions (Wang et al., 2012; Joshi and Gold, 2020; Wang and Munoz, 2021; Burlingham et al., 2022; Wang and Munoz, 2024), but also by regions that are less involved in eye-movement regulation, such as the locus coeruleus (Mathôt, 2018; Joshi and Gold, 2020; Strauch et al., 2022).

Changes the pupil size are associated with changes in arousal, which in turn is driven by locus coeruleus function (Bradley et al., 2008; Mathôt, 2018; Wang et al., 2018; Ross and Van Bockstaele, 2021; Burlingham et al., 2022). The pupil size is thought to index listening effort via the arousal system, such that the higher cognitive load associated with effort increases arousal, which, in turn, increases the pupil size (Winn et al., 2018; Zekveld et al., 2018; Fink et al., 2023). Changes in eye movements are not tied into the arousal system and may thus involve different processes. Small eye movements (microsaccades) occur during fixation that would largely be considered involuntary (Martinez-Conde et al., 2009; Martinez-Conde et al., 2013; Alexander and Martinez-Conde, 2019; although microsaccades can also be voluntarily initiated under rare conditions: Willeke et al., 2019). Eye-movement reductions due to speech masking have been observed while participants fixate on a point (Contadini-Wright et al., 2023; Cui and Herrmann, 2023), suggesting a role of involuntary eye movements. This is also consistent with work showing reductions in eye movements during fixation under high compared to low memory load (Dalmaso et al., 2017; Kadosh et al., 2024). However, a listener can also voluntarily change their eye movements (Pierce et al., 2019), for example, by fixating on a specific location to avoid visual distractions rather than engaging in visual search under challenging listening conditions.

That changes in pupil size and eye movements may index listening effort differently is also suggested by the absence of a between-participant correlation (although saccade rate negatively correlated with pupil area when limited to the first block) and by recent work on disengagement from effortful listening (Herrmann and Ryan, 2024). When listeners disengage from listening (because comprehension is impossible) and listening effort is low as a result, the pupil area decreases (indexing lower arousal), whereas eye movements are also reduced, which would indicate greater listening effort (Herrmann and Ryan, 2024). The reasons for this are not fully understood. It is possible that when individuals disengage and orient their attention mentally inwards (e.g., think about their day) they reduce their eye movements (Herrmann and Ryan, 2024), but this requires further investigation. These previous data and to some extent the current data suggest, nevertheless, that eye movements might index effortful listening differently than the pupil size, possibly pointing to an advantage of capitalizing on both measures when possible (cf. Contadini-Wright et al., 2023).

## Conclusions

The current study investigated whether eye movements are sensitive to listening effort induced through speech masking during continuous story listening, whether changes in eye movements with increased speech masking depend on visual-stimulation conditions, and whether eye-movement changes differ between younger and older adults. The results show that eye movements decrease (as indicated by fixation duration, gaze dispersion, and saccade rate) as speech masking and associated listening effort increase, and that this eye-movement reduction appears to be independent of visual-stimulation conditions and does not differ between younger and older adults. The data suggest that eye movements could potentially be used as a measure of listening effort during continuous story listening.

## Data Availability

Data will be available to other researchers upon reasonable request to the corresponding author.

## Author Contributions

BH: Conceptualization, methodology, validation, formal analysis, investigation, data curation, writing - original draft, writing - review & editing, visualization, supervision, project management, funding acquisition. FS: Methodology, formal analysis, writing - review & editing. AW: Methodology, formal analysis, writing - review & editing.

## Acknowledgements

The research was supported by the Canada Research Chair program (CRC-2023-00383), the Natural Sciences and Engineering Research Council of Canada (NSERC Discovery Grant: RGPIN-2021-02602), the Canadian Institutes of Health Research (CIHR: 186236). We thank Chistie Tsagopoulos for her help during data collection.

## Statements and Declarations

The author has no conflicts or competing interests.

## References

Alexander RG, Martinez-Conde S (2019) Fixational Eye Movements. In: Eye Movement Research: An Introduction to its Scientific Foundations and Applications (Klein C, Ettinger U, eds), pp 73–115. Cham: Springer International Publishing.

Alvarez GA, Franconeri SL (2007) How many objects can you track?: Evidence for a resource-limited attentive tracking mechanism. Journal of Vision 7:1–10.

Ayasse ND, Wingfield A (2018) A Tipping Point in Listening Effort: Effects of Linguistic Complexity and Age-Related Hearing Loss on Sentence Comprehension. Trends in Hearing 22:2331216518790907.

Benwell CSY, Keitel C, Harvey M, Gross J, Thut G (2018) Trial-by-trial co-variation of pre-stimulus EEG alpha power and visuospatial bias reflects a mixture of stochastic and deterministic effects. European Journal of Neuroscience 48:2566–2584.

Bilger RC (1984) Manual for the clinical use of the revised SPIN Test. Champaign, IL, USA: The University of Illinois.

Borghini G, Hazan V (2018) Listening Effort During Sentence Processing Is Increased for Non-native Listeners: A Pupillometry Study. Frontiers in Neuroscience 12.

Bradley MM, Miccoli L, Escrig MA, Lang PJ (2008) The pupil as a measure of emotional arousal and autonomic activation. Psychophysiology 45:602–607.

Brisson J, Mainville M, Mailloux D, Beaulieu C, Serres J, Sirois S (2013) Pupil diameter measurement errors as a function of gaze direction in corneal reflection eyetrackers. Behavior Research Methods 45:1322–1331.

Brodbeck C, Simon JZ (2020) Continuous speech processing. Current Opinion in Physiology 18:25–31.

Broderick MP, Anderson AJ, Di Liberto GM, Crosse MJ, Lalor EC (2018) Electrophysiological Correlates of Semantic Dissimilarity Reflect the Comprehension of Natural, Narrative Speech. Current Biology 28:803–809.

Burlingham CS, Mirbagheri S, Heeger DJ (2022) A unified model of the task-evoked pupil response. Science Advances 8:eabi9979.

Cavanagh P, Alvarez GA (2005) Tracking multiple targets with multifocal attention. Trends in Cognitive Sciences 9:349–354.

Cohen J (1988) Statistical power analysis for the behavioral sciences, 2nd ed Edition. Hillsdale, N.J: L. Erlbaum Associates.

Contadini-Wright C, Magami K, Mehta N, Chait M (2023) Pupil dilation and microsaccades provide complementary insights into the dynamics of arousal and instantaneous attention during effortful listening. The Journal of Neuroscience 43:4856–4866.

Cruickshanks KJ, Wiley TL, Tweed TS, Klein BEK, Klein R, Mares-Perlman JA, Nondahl DM (1998) Prevalence of Hearing Loss in Older Adults in Beaver Dam, Wisconsin. American Journal of Epidemiology 148:879–886.

Cui ME, Herrmann B (2023) Eye Movements Decrease during Effortful Speech Listening. The Journal of Neuroscience 43:5856–5869.

Dalmaso M, Castelli L, Scatturin P, Galfano G (2017) Working memory load modulates microsaccadic rate. Journal of Vision 17:6.

Ding N, Simon JZ (2014) Cortical entrainment to continuous speech: functional roles and interpretations. Frontiers in Human Neuroscience 8.

Eckert MA, Teubner-Rhodes S, Vaden Jr. KI (2016) Is Listening in Noise Worth It? The Neurobiology of Speech Recognition in Challenging Listening Conditions. Ear & Hearing 37:101S–110S.

Engbert R (2006) A microcosm for research on oculomotor control, attention, and visual perception. Progress in Brain Research 154:177–192.

Engbert R, Kliegl R (2003) Microsaccades uncover the orientation of covert attention. Vision Research 43:1035–1045.

Farahani M, Parsa V, Herrmann B, Kadem M, Johnsrude IS, Doyle PC (2020) An Auditory-Perceptual and Pupillometric Study of Vocal Strain and Listening Eort in Adductor Spasmodic Dysphonia. Applied Sciences 10:5907.

Ferguson SH, Jongman A, Sereno JA, Keum KA (2010) Intelligibility of Foreign-Accented Speech for Older Adults with and without Hearing Loss. Journal of the American Academy of Audiology 21:153–162.

Fiedler L, Seifi Ala T, Graversen C, Alickovic E, Lunner T, Wendt D (2021) Hearing Aid Noise Reduction Lowers the Sustained Listening Effort During Continuous Speech in Noise—A Combined Pupillometry and EEG Study. Ear Hear 42.

Fink L, Simola J, Tavano A, Lange E, Wallot S, Laeng B (2023) From pre-processing to advanced dynamic modeling of pupil data. Behavior Research Methods.

Goman AM, Lin FR (2016) Prevalence of Hearing Loss by Severity in the United States. American Journal of Public Health 106:1820–1822.

Gordon MS, Daneman M, Schneider BA (2009) Comprehension of Speeded Discourse by Younger and Older Listeners. Experimental Aging Research 35:277–296.

Hayes TR, Petrov AA (2016) Mapping and correcting the influence of gaze position on pupil size measurements. Behavior Research Methods 48:510–527.

He X, Raghavan VS, Mesgarani N (2024) Distinct roles of SNR, speech Intelligibility, and attentional effort on neural speech tracking in noise. BioRxiv.

Helfer KS, Freyman RL (2008) Aging and Speech-on-Speech Masking. Ear & Hearing 29:87–98.

Helfer KS, Jesse A (2021) Hearing and speech processing in midlife. Hearing Research 402:108097.

Herrmann B, Johnsrude IS (2018a) Neural signatures of the processing of temporal patterns in sound. The Journal of Neuroscience 38:5466–5477.

Herrmann B, Johnsrude IS (2018b) Attentional State Modulates the Effect of an Irrelevant Stimulus Dimension on Perception. Journal of Experimental Psychology: Human Perception and Performance 44:89–105.

Herrmann B, Johnsrude IS (2020a) A Model of Listening Engagement (MoLE). Hearing Research 397:108016.

Herrmann B, Johnsrude IS (2020b) Absorption and enjoyment during listening to acoustically masked stories. Trends in Hearing 24:1–18.

Herrmann B, Ryan JD (2024) Pupil size and eye movements differently index effort in both younger and older adults. BioRxiv.

Herrmann B, Maess B, Johnsrude IS (2018) Aging Affects Adaptation to Sound-Level Statistics in Human Auditory Cortex. The Journal of Neuroscience 38:1989–1999.

Herrmann B, Maess B, Johnsrude IS (2023) Sustained responses and neural synchronization to amplitude and frequency modulation in sound change with age. Hearing Research 428:108677.

Holmes AP, Friston KJ (1998) Generalisability, Random Effects & Population Inference. NeuroImage 7:S754.

Humes LE (2019) Examining the Validity of the World Health Organization’s Long-Standing Hearing Impairment Grading System for Unaided Communication in Age-Related Hearing Loss. American Journal of Audiology 28:810–818.

Irsik VC, Johnsrude IS, Herrmann B (2022a) Age-related deficits in dip-listening evident for isolated sentences but not for spoken stories. Scientific Reports 12:5898.

Irsik VC, Johnsrude IS, Herrmann B (2022b) Neural activity during story listening is synchronized across individuals despite acoustic masking. Journal of Cognitive Neuroscience 34:933–950.

Johansson R, Holsanova J, Holmqvist K (2006) Pictures and Spoken Descriptions Elicit Similar Eye Movements During Mental Imagery, Both in Light and in Complete Darkness. Cognitive Science 30:1053–1079.

Johansson R, Holsanova J, Homqvist K (2011) The Dispersion of Eye Movements During Visual Imagery is Related to Individual Differences in Spatial Imagery Ability. In: Annual Meeting of the Cognitive Science Society, pp 1200–1205: Proceedings of the Annual Meeting of the Cognitive Science Society.

Johansson R, Holsanova J, Dewhurst R, Holmqvist K (2012) Eye movements during scene recollection have a functional role, but they are not reinstatements of those produced during encoding. Journal of Experimental Psychology: Human Perception and Performance 38:1289–1314.

Joshi S, Gold JI (2020) Pupil Size as a Window on Neural Substrates of Cognition. Trends in Cognitive Sciences 24:466–480.

Kadem M, Herrmann B, Rodd JM, Johnsrude IS (2020) Pupil dilation is sensitive to semantic ambiguity and noise. Trends in Hearing 24:1–16.

Kadosh O, Inbal K, Snir H, Bonneh YS (2024) Oculomotor inhibition markers of working memory load. Scientific Reports 14:1872.

Kılıç S, Sendesen E, Aslan F, Erbil N, Aydın Ö, Türkyılmaz D (2024) Investigating Sensitivity to Auditory Cognition in Listening Effort Assessments: A Simultaneous EEG and Pupillometry Study. Brain and Behavior 14:e70135.

Knapen T, de Gee JW, Brascamp J, Nuiten S, Hoppenbrouwers S, Theeuwes J (2016) Cognitive and Ocular Factors Jointly Determine Pupil Responses under Equiluminance. PLoS ONE 11:e0155574.

Koelewijn T, Zekveld AA, Festen JM, Kramer SE (2014) The influence of informational masking on speech perception and pupil response in adults with hearing impairment. The Journal of the Acoustical Society of America 135:1596–1606.

Koelewijn T, Zekveld AA, Lunner T, Kramer SE (2018) The effect of reward on listening effort as reflected by the pupil dilation response. Hearing Research 367:106–112.

Kosch T, Hassib M, Woźniak PW, Buschek D, Alt F (2018) Your Eyes Tell: Leveraging Smooth Pursuit for Assessing Cognitive Workload. In: CHI Conference on Human Factors in Computing Systems, p Paper 436. Montreal, QC, Canada.

Kuchinsky SE, Ahlstrom JB, Vaden Jr. KI, Cute SL, Humes LE, Dubno JR, Eckert MA (2013) Pupil size varies with word listening and response selection difficulty in older adults with hearing loss. Psychophysiology 50:23–34.

Lalor EC, Foxe JJ (2010) Neural responses to uninterrupted natural speech can be extracted with precise temporal resolution. European Journal of Neuroscience 31:189–193.

Lindquist MA (2008) The Statistical Analysis of fMRI Data. Statistical Science 23:439–464.

Liu Z-X, Shen K, Olsen RK, Ryan JD (2018) Age-related changes in the relationship between visual exploration and hippocampal activity. Neuropsychologia 119:81–91.

Martinez-Conde S, Otero-Millan J, Macknik SL (2013) The impact of microsaccades on vision: towards a unified theory of saccadic function. Nature Reviews Neuroscience 14:83–96.

Martinez-Conde S, Macknik SL, Troncoso XG, Hubel DH (2009) Microsaccades: a neurophysiological analysis. Trends in Neurosciences 32:463–475.

Mathiesen SL, Van Hedger SC, Irsik VC, Bain MM, Johnsrude IS, Herrmann B (2024) Exploring age differences in absorption and enjoyment during story listening. Psychology International 6:667–684.

Mathôt S (2018) Pupillometry: Psychology, physiology, and function. Journal of Cognition 1.

Mazloum-Farzaghi N, Shing N, Mendoza L, Barense MD, Ryan JD, Olsen RK (2022) The impact of aging and repetition on eye movements and recognition memory. Aging, Neuropsychology, and Cognition:1–27.

McIntire LK, McIntire JP, McKinley RA, Goodyear C (2014) Detection of vigilance performance with pupillometry. In: Symposium on Eye Tracking Research and Applications, pp 167–174. Safety Harbor, Florida, USA: ACM.

McLaughlin DJ, Zink ME, Gaunt L, Reilly J, Sommers MS, Van Engen KJ, Peelle JE (2023) Give me a break! Unavoidable fatigue effects in cognitive pupillometry. Psychophysiology 60:e14256.

Moore BCJ (2007) Cochlear Hearing Loss: Physiological, Psychological and Technical Issues. West Sussex, Engand: John Wiley & Sons, Ltd.

Nakayama M, Takahashi K, Shimizu Y (2002) The act of task difficulty and eye-movement frequency for the ‘Oculo-motor indices’. In, pp 37–42. New York, NY, USA: ACM.

Neagu M-B, Kressner AA, Relaño-Iborra H, Bækgaard P, Dau T, Wendt D (2023) Investigating the Reliability of Pupillometry as a Measure of Individualized Listening Effort. Trends in Hearing 27:23312165231153288.

Ohlenforst B, Zekveld AA, Lunner T, Wendt D, Naylor G, Wang Y, Versfeld NJ, Kramer SE (2017) Impact of stimulus-related factors and hearing impairment on listening effort as indicated by pupil dilation. Hearing Research 351:68–79.

Otero-Millan J, Troncoso XG, Macknik SL, Serrano-Pedraza I, Martinez-Conde S (2008) Saccades and microsaccades during visual fixation, exploration, and search: Foundations for a common saccadic generator. Journal of Vision 8:21–21.

Oxenham AJ, Bacon SP (2003) Cochlear Compression: Perceptual Measures and Implications for Normal and Impaired Hearing. Ear & Hearing 24:352–366.

Pan J, Sun X, Park E, Kaufmann M, Klimova M, McGuire JT, Ling S (2024) The effects of emotional arousal on pupil size depend on luminance. Scientific Reports 14:21895.

Pandey PR, Herrmann B (in press) The influence of semantic context on the intelligibility benefit from speech glimpses in younger and older adults. Journal of Speech, Language, and Hearing Research.

Panela RA, Copelli F, Herrmann B (2024) Reliability and generalizability of neural speech tracking in younger and older adults. Neurobiology of Aging 134:165–180.

Payne BR, Silcox JW (2019) Aging, context processing, and comprehension. In: Psychology of Learning and Motivation (Federmeier KD, ed), pp 215–264: Academic Press.

Peelle JE (2018) Listening Effort: How the Cognitive Consequences of Acoustic Challenge Are Reflected in Brain and Behavior. Ear & Hearing 39:204–214.

Penny WD, Holmes AJ (2007) Random Effects Analysis. In: Statistical Parametric Mapping (Friston K, Ashburner J, Kiebel S, Nichols T, Penny W, eds), pp 156–165. London: Academic Press.

Pichora-Fuller MK, Levitt H (2012) Speech Comprehension Training and Auditory and Cognitive Processing in Older Adults. American Journal of Audiology 21:351–357.

Pichora-Fuller MK, Kramer SE, Eckert MA, Edwards B, Hornsby BWY, Humes LE, Lemke U, Lunner T, Matthen M, Mackersie CL, Naylor G, Phillips NA, Richter M, Rudner M, Sommers MS, Tremblay KL, Wingfield A (2016) Hearing Impairment and Cognitive Energy: The Framework for Understanding Effortful Listening (FUEL). Ear & Hearing 37 Suppl 1:5S–27S.

Pierce JE, Clementz BA, McDowell JE (2019) Saccades: Fundamentals and Neural Mechanisms. In: Eye Movement Research: An Introduction to its Scientific Foundations and Applications (Klein C, Ettinger U, eds), pp 11–71. Cham: Springer International Publishing.

Pierrot-Deseilligny C, Milea D, Müri RM (2004) Eye movement control by the cerebral cortex. Current Opinion in Neurology 17.

Plack CJ (2014) The sense of hearing. New York, USA: Psychology Press.

Presacco A, Simon JZ, Anderson S (2016) Evidence of degraded representation of speech in noise, in the aging midbrain and cortex. Journal of Neurophysiology 116:2346–2355.

Presacco A, Simon JZ, Anderson S (2019) Speech-in-noise representation in the aging midbrain and cortex: Effects of hearing loss. PLoS ONE 14:e0213899.

Ross JA, Van Bockstaele EJ (2021) The Locus Coeruleus-Norepinephrine System in Stress and Arousal: Unraveling Historical, Current, and Future Perspectives. Frontiers in Psychiatry 11.

Ryan JD, Leung G, Turk-Browne NB, Hasher L (2007) Assessment of age-related changes in inhibition and binding using eye movement monitoring. Psychology and Aging 22:239–250.

Scholl BJ (2009) What have we learned about attention from multiple object tracking (and vice versa)? In: Computation, cognition, and Pylyshyn (Dedrick D, Trick L, eds), pp 49–78. Cambridge, MA, USA: MIT Press.

Seifi Ala T, Graversen C, Wendt D, Alickovic E, Whitmer WM, Lunner T (2020) An exploratory Study of EEG Alpha Oscillation and Pupil Dilation in Hearing-Aid Users During Effortful listening to Continuous Speech. PLOS ONE 15:e0235782.

Sobon KA, Taleb NM, Buss E, Grose JH, Calandruccio L (2019) Psychometric function slope for speech-in-noise and speech-in-speech: Effects of development and aging. The Journal of the Acoustical Society of America 145:EL284-EL290.

Sparks DL (2002) The brainstem control of saccadic eye movements. Nature Reviews Neuroscience 3:952–964.

Stevens G, Flaxman S, Brunskill E, Mascarenhas M, Mathers CD, Finucane M, on behalf of the Global Burden of Disease Hearing Loss Expert G (2013) Global and regional hearing impairment prevalence: an analysis of 42 studies in 29 countries. European Journal of Public Health 23:146–152.

Strauch C, Wang C-A, Einhäuser W, Van der Stigchel S, Naber M (2022) Pupillometry as an integrated readout of distinct attentional networks. Trends in Neurosciences 45:635–647.

Suzuki Y, Minami T, Laeng B, Nakauchi S (2019) Colorful glares: Effects of colors on brightness illusions measured with pupillometry. Acta Psychologica 198:102882.

Thurman SM, Cohen Hoffing RA, Madison A, Ries AJ, Gordon SM, Touryan J (2021) “Blue Sky Effect”: Contextual Influences on Pupil Size During Naturalistic Visual Search. Frontiers in Psychology 12.

Unsworth N, Robison MK (2016) Pupillary correlates of lapses of sustained attention. Cognitive, Affective, & Behavioral Neuroscience 16:601–615.

van der Wel P, van Steenbergen H (2018) Pupil dilation as an index of effort in cognitive control tasks: A review. Psychonomic Bulletin & Review 25:2005–2015.

Walter K, Bex P (2021) Cognitive load influences oculomotor behavior in natural scenes. Scientific Reports 11:12405.

Wang C-A, Munoz DP (2021) Coordination of Pupil and Saccade Responses by the Superior Colliculus. Journal of Cognitive Neuroscience 33:919–932.

Wang C-A, Munoz DP (2024) Linking the Superior Colliculus to Pupil Modulation. In: Modern Pupillometry: Cognition, Neuroscience, and Practical Applications (Papesh MH, Goldinger SD, eds), pp 77–98. Cham: Springer International Publishing.

Wang C-A, Boehnke SE, White BJ, Munoz DP (2012) Microstimulation of the Monkey Superior Colliculus Induces Pupil Dilation Without Evoking Saccades. The Journal of Neuroscience 32:3629.

Wang C-A, Baird T, Huang J, Coutinho JD, Brien DC, Munoz DP (2018) Arousal Effects on Pupil Size, Heart Rate, and Skin Conductance in an Emotional Face Task. Frontiers in Neurology 9.

Wendt D, Dau T, Hjortkjær J (2016) Impact of Background Noise and Sentence Complexity on Processing Demands during Sentence Comprehension. Frontiers in Psychology 7:Article 345.

Wendt D, Hietkamp RK, Lunner T (2017) Impact of Noise and Noise Reduction on Processing Effort: A Pupillometry Study. Ear & Hearing 38:690–700.

WHO (2024) Deafness. In. https://www.who.int/news-room/fact-sheets/detail/deafness-and-hearing-loss.

Widmann A, Engbert R, Schröger E (2014) Microsaccadic Responses Indicate Fast Categorization of Sounds: A Novel Approach to Study Auditory Cognition. The Journal of Neuroscience 34:11152–11158.

Widmann A, Herrmann B, Scharf F (2025) Pupillometry is sensitive to speech masking during story listening: Acommentary on the critical role of modeling temporal trends. Journal of Neuroscience Methods 413:110299.

Willeke KF, Tian X, Buonocore A, Bellet J, Ramirez-Cardenas A, Hafed ZM (2019) Memory-guided microsaccades. Nature Communications 10:3710.

Wilson RH, McArdle RA, Watts KL, Smith SL (2012) The Revised Speech Perception in Noise Test (R-SPIN)in a Multiple Signal-to-Noise Ratio Paradigm. Journal of the American Academy of Audiology 23:590–605.

Winn MB (2016) Rapid Release From Listening Effort Resulting From Semantic Context, and Effects of Spectral Degradation and Cochlear Implants. Trends in Hearing 20:2331216516669723.

Winn MB, Teece KH (2021) Listening Effort Is Not the Same as Speech Intelligibility Score. Trends in Hearing 25:1–26.

Winn MB, Edwards JR, Litovsky RY (2015) The Impact of Auditory Spectral Resolution on Listening Effort Revealed by Pupil Dilation. Ear & Hearing 36:e153–e165.

Winn MB, Wendt D, Koelewijn T, Kuchinsky SE (2018) Best Practices and Advice for Using Pupillometry to Measure Listening Effort: An Introduction for Those Who Want to Get Started. Trends in Hearing 22:2331216518800869.

Worsley KJ, Liao CH, Aston J, Petre V, Duncan GH, Morales F, Evans AC (2002) A General Statistical Analysis for fMRI Data. NeuroImage 15:1–15.

Zekveld AA, Kramer SE (2014) Cognitive processing load across a wide range of listening conditions: Insights from pupillometry. Psychophysiology 51:277–284.

Zekveld AA, Kramer SE, Festen JM (2010) Pupil response as an indication of effortful listening: the influence of sentence intelligibility. Ear & Hearing 31.

Zekveld AA, Koelewijn T, Kramer SE (2018) The Pupil Dilation Response to Auditory Stimuli: Current State of Knowledge. Trends in Hearing 22:1–25.

Zekveld AA, Heslenfeld DJ, Johnsrude IS, Versfeld NJ, Kramer SE (2014) The eye as a window to the listening brain: Neural correlates of pupil size as a measure of cognitive listening load. NeuroImage 101:76–86.

Zekveld AA, van Scheepen JAM, Versfeld NJ, Veerman ECI, Kramer SE (2019) Please try harder! The influence of hearing status and evaluative feedback during listening on the pupil dilation response, salivacortisol and saliva alpha-amylase levels. Hearing Research 381:107768.

Zekveld Adriana A, Pielage H, Versfeld Niek J, Kramer Sophia E (2023) The Influence of Hearing Loss on the Pupil Response to Degraded Speech. Journal of Speech, Language, and Hearing Research 66:4083–4099.

Zhang Y, Malaval F, Lehmann A, Deroche MLD (2022) Luminance effects on pupil dilation in speech-in-noise recognition. PLOS ONE 17:e0278506.

Zhao S, Bury G, Milne A, Chait M (2019) Pupillometry as an Objective Measure of Sustained Attention in Young and Older Listeners. Trends in Hearing 23:1–21.

